# Extensive Decoupling of Metabolic Genes in Cancer

**DOI:** 10.1101/008946

**Authors:** Ed Reznik, Chris Sander

## Abstract

Tumorigenesis involves, among other factors, the alteration of metabolic gene expression to support malignant, unrestrained proliferation. Here, we examine how the altered metabolism of cancer cells is reflected in changes in co-expression patterns of metabolic genes between normal and tumor tissues. Our emphasis on changes in the interactions of pairs of genes, rather than on the expression levels of individual genes, exposes changes in the activity of metabolic pathways which do not necessarily show clear patterns of over- or under-expression. We report the existence of key metabolic genes which act as hubs of differential co-expression, showing significantly different co-regulation patterns between normal and tumor states. Notably, we find that the extent of differential co-expression of a gene is only weakly correlated with its differential expression, suggesting that the two measures probe different features of metabolism. By leveraging our findings against existing pathway knowledge, we extract networks of functionally connected differentially co-expressed genes and the transcription factors which regulate them. Doing so, we identify a previously unreported network of dysregulated metabolic genes in clear cell renal cell carcinoma transcriptionally controlled by the transcription factor HNF4A. While HNF4A shows no significant differential expression, the co-expression HNF4A and several of its regulated target genes in normal tissue is completely abrogated in tumor tissue. Finally, we aggregate the results of differential co-expression analysis across seven distinct cancer types to identify pairs of metabolic genes which may be recurrently dysregulated. Among our results is a cluster of four genes, all located in the mitochondrial electron transport chain, which show significant loss of co-expression in tumor tissue, pointing to potential mitochondrial dysfunction in these tumor types.

## Introduction

All cellular events, from the transduction of signals to the translation of nucleic acids, rely on the interaction of molecular entities. Indeed, one may argue that the fundamental unit of a biological network is not its constituent components (*e.g.* proteins or genes), but rather the edges representing the interactions between them. Then, it follows that the manifestation of disease, of a deranged phenotype of this network, should be evident by observing changes in the wiring and activity of these edges.

Here, we study the interactions between pairs of genes encoding metabolic enzymes, and how these interactions change in the course of transformation of normal cells to malignant tumor. This notion of studying “interactions” is particularly important for understanding the network of coupled enzymatic reactions which constitute metabolism. It is well-known that tumors, which are under strong selection for proliferative capacity, must re-organize their metabolism in order to deliver the precursors and energy needed to grow as quickly as possible. Otto Warburg published a series of key findings highlighting a fundamental dysregulation in glycolytic metabolism in cancer, whereby cancer cells metabolized high levels of glucose to lactate [28]. Some of the earliest chemotherapies (*e.g.* methotrexate) targeted a metabolic phenotype which distinguished tumor from normal tissue. In recent years, an invigorated field has identified a number of distinct “metabolic lesions” in various tumors, including, for example, the preferential expression of PKM2 [48] and the presence of an oncometabolite, 2-hydroxyglutarate, in cells with activating IDH1 and IDH2 mutations [34].

Our use of the term “interaction” above is loose: for the purposes of our study, which focuses on the analysis of gene expression data, we say that two metabolic genes putatively interact if we observe they are co-expressed. This co-expression may occur by chance, or as a result of co-regulation by a set of common factors. Furthermore, while strong co-expression is more likely to occur between proteins which physically interact with each other, the highly networked structure of the metabolic network suggests that even genes residing in opposing corners of metabolism may be fundamentally coupled to each other. Regardless of the source of co-expression, our goal is to identify regions of the metabolic network whose co-expression patterns appear fundamentally different between normal and cancerous tissue samples. Put another way, we intentionally search for cases where two genes are co-expressed in one manner in normal tissue, and then co-expressed in an entirely different manner in the tumor tissue. Our approach follows other studies employing techniques to detect so-called “differential co-expression” of genes [9, 10, 17, 25, 26, 33, 41, 46].

Differential expression analysis is the standard method for identifying comparing the expression patterns of genes across conditions. Aside from its ubiquitous use in research, several large-scale surveys of differential expression focusing exclusively on metabolic genes in cancer have been completed [23, 36]. In contrast, while a handful of publications have examined differential co-expression in various cancer settings (for example, [2, 6, 7, 10, 29]), differential co-expression analysis remains largely absent in most studies of gene expression and (to our knowledge), no survey of differential co-expression among metabolic genes in cancer ahs been undertaken. This is, at least in part, due to the requirement for large sample sizes in order to detect statistically significant differential co-expression patterns. Here, we embark on such a large-scale analysis of RNA-Seq data from 3000 samples of primary tumor and adjacent normal samples from seven distinct tissues, and focus our attention squarely on the expression patterns of 1789 metabolic genes. Among our main findings is the (previously known, see [24], but potentially under-appreciated) observation that genes with strong differential co-expression patterns are not necessarily differential expressed. A relatively large fraction of the genes we identify in our study show no substantial difference in their absolute expression between tumor and normal tissue, but nevertheless exhibit recurrent differential co-expression.

The orthogonality of differential expression and differential co-expression described above suggests that, to detect changes in the activity of a pathway, one must separately investigate the unilateral increase/decrease of enzyme levels, as well changes in their coordinated co-expression. In the first case, the expression of a large set of genes (for example, those in a long, linear metabolic pathway) may be synchronously upregulated. This coordinated up-regulation of transcription may, for example, enable the pathway to carry substantially more metabolic flux. In the second, perhaps more subtle case, the characteristic pattern of flux through a pathway may be re-wired (as illustrated in Figure 1). In Figure 1, the mechanism for this re-wiring is transcriptional, but in principle this type of coupling may arise through a variety of distinct mechanisms (such, as, for example, post-translational modification). In both cases, changes in intra- or extra-cellular conditions across a set of samples induces variation in the expression of genes. However, the manifestation of these changes may be hidden from either differential expression or differential co-expression analysis. Thus, we will repeatedly argue that both differential expression and differential co-expression analysis should play central, complementary roles in the analysis of gene expression data [25].

**Figure 1.**
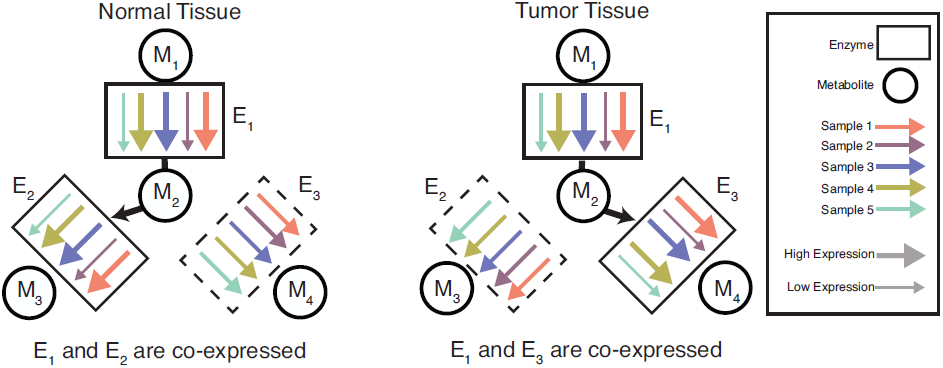
Differential co-expression can signal a change in the activity of a pathway. Each arrow represents the level of expression of an enzymatic gene from a single sample (*e.g.* a patient, so that all arrows of the same color derive from the same sample). In normal tissue **(A)**, the expression of genes encoding enzymes *E*1 and *E*2 are strongly correlated, and the expression of *E*_1_ and *E*_3_ are uncorrelated. In tumor tissue **(B)**, the expression of genes encoding enzymes *E*_1_ and *E*_3_ are strongly correlated, and the expression of *E*_1_ and *E*_2_ are uncorrelated. If we assume that enzyme activity is correlated with expression, then we may hypothesize that the metabolic flux exiting from *E*_1_ is coupled to flux in *E*_2_ in normal tissue, and to flux in *E*_3_ in tumor tissue. Note that the average expression of all enzymes remains constant between tumor and normal conditions, so that a differential expression analysis would be unlikely to identify the expression of these genes as anamolous.

The results to be presented will encompass a variety of analyses, studying differential co-expression patterns first across two cancer types for which we have the most data available (breast and clear cell renal cell carcinomas, KIRC), and then expanding to include five other cancer types (lung, thyroid, prostate, liver, and head and neck), as described in Table 1. In the course of doing so, we propose two simple, but novel, analyses which integrate pathway information to assess the functional role of differentially co-expressed gene pairs. We examine the association between differential co-expression and differential expression, and identify genes which are strongly enriched for one measure but not the other. By leveraging our findings against regulatory (*i.e.* transcription factor binding) data, we identify transcription factors whose targets are highly enriched for differential co-expression. Among our findings is a previously unreported loss of co-expression between HNF4A, a transcription factor, and its regulatory targets in KIRC. Finally, we leverage the scale of our study to complete a “Pan-Cancer” analysis of differential co-expression, searching for those pairs of metabolic genes which are recurrently differentially co-expressed across multiple cancer types. Our results highlight a small group of four mitochondrial electron transport chain (ETC) genes which are recurrently differentially co-expressed, hinting at a fundamental alteration in the function of the ETC in tumors.

**Table 1.** Cancer data used in this study.

| Study | Number Tu-<br>mor Samples | Number<br>Normal<br>Samples | Individual<br>Analysis | PanCan<br>Analysis |
| --- | --- | --- | --- | --- |
| Breast Cancer | 914 | 106 | **X** | **X** |
| Clear Cell Renal Cell Car-<br>cinoma | 480 | 71 | **X** | **X** |
| Head and Neck Squamous<br>Cell Carcinoma | 303 | 37 |  | **X** |
| Liver Cancer | 103 | 49 |  | **X** |
| Lung Adenocarcinoma | 446 | 57 |  | **X** |
| Prostate Adenocarcinoma | 176 | 44 |  | **X** |
| Thyroid Cancer | 482 | 58 |  | **X** |
| **Total** | **2904** | **422** |  |  |

## Results

### Calculation of Changes in Gene Co-Expression

We begin by describing the methodology, broadly illustrated in Figure 2, to detect changes in co-expression patterns between normal and tumor samples. After obtaining RNA-Seq data, we calculate the Spearman correlation (a non-parametric measure of the correlation of two random variables employing ranks) of each pair of genes *i, j*, and record the p-value *p*_*ij*_ associated with this correlation. These calculations are performed separately for tumor and normal samples. To account for multiple hypothesis testing, we apply the conservative Bonferonni correction [16], yielding corresponding adjusted p-values *p*̂. The results of these correlation calculations are stored in two matrices, *C*^*T*^ and *C*^*N*^ (corresponding to tumor and normal samples, respectively), with entries

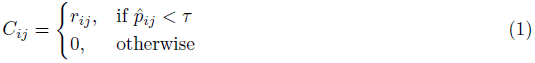

**Figure 2.**
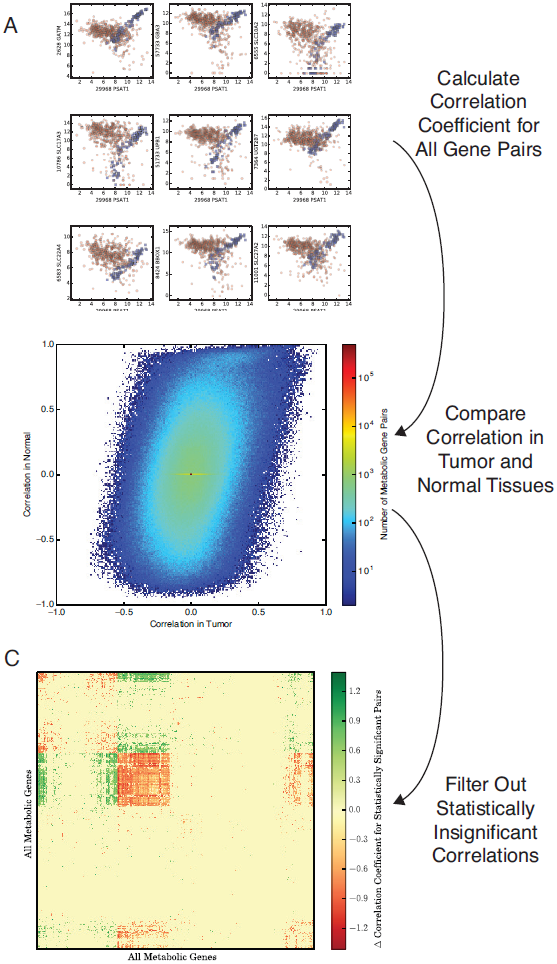
Outline of the method to calculate differential co-expression. (**A**) Calculate the co-expression for each pair of metabolic genes across tumor (red) and normal (blue) samples, respectively. (**B**) For each pair of genes in a given tumor type (*e.g.* breast), compare the Spearman correlation coefficient in tumor and normal samples. Most pairs of genes show very similar co-expression in both tumor and normal samples (reflected in the high density of points in the center of the plot). More rarely, a pair of genes will show significantly different co-expression between normal and tumor samples (*e.g.* bottom right and top left corners). (**C**) Using the statistical methodology detailed in 4, filter out insiginificant differences in correlation coefficients. Retain the remaining (significant) differences in correlations in the matrix **D**. The filtered results can then be analyzed further to identify regions of metabolism enriched for differential co-expression.

Here, *τ* is a significance threshold for our Bonferroni-corrected p-values. Throughout the manuscript unless otherwise stated, we employ a threshold *τ* = 1×10^*−*2^.

Our goal is to identify significant differences between the strength of co-expression (as quantified by the correlation coefficients) in tumor and normal samples. Such a comparison of sample correlation coefficients must be done with care. In fact, the difference between two correlation coefficients is not sufficient information to determine how often such a difference would appear by chance. We offer an example to illustrate this phenomenon. Very small correlation coefficients (say, *r*_1_ = 0.1, *r*_2_ = –0.1) may appear in random, uncorrelated data simply by chance. In this case, the difference between the two correlation coefficients (*r*_1_ *− r*_2_ = 0.2) should be categorized as statistically insignificant because it is quite likely to happen by chance. On the other hand, the same difference for two very large correlation coefficients (say, *r*_1_ = 0.99, *r*_2_ = 0.79) appears less likely to happen by chance; instead, this difference is more likely to arise via the corruption of a nearly perfect correlation by a confounding factor or noise.

The basis of this intuition is that very large correlation coefficients are observed quite rarely by chance. More importantly, the variance of the correlation coefficient estimated from the data (referred to as the *sample* correlation coefficient, *r*) depends on the value of the *true* correlation coefficient underlying the data (referred to as the *population* correlation coefficient, *ρ*). In particular, the variance of sample correlation coefficient is approximately

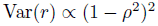

Thus, as the population correlation coefficient tends to ±1, the variance of the sample correlation coefficient asymptotically approaches zero. This dependence of the variance of *r* on *ρ* itself makes it very difficult to carry out hypothesis tests comparing two sample correlation coefficients. A standard method for testing for a difference between correlation coefficients is to employ a transformation to *stabilize* the variances, making them independent of *ρ*. Here, we use the Fisher r to z transformation:

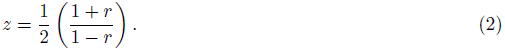

The change of variables in Eq. (2) is well-known, and has been used in prior work on differential co-expression [9]. When applied to data drawn from a bivariate normal distribution, this transformation yields a quantity which is approximately normally distributed with variance 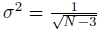 independent of the population mean, with *N* equal to the size of the population. By applying this transformation to our measured correlation coefficients in normal tissue and tumor samples, we are able to apply a Z-test to determine if the correlation coefficients 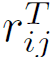 and 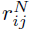 are significantly different. In particular, the quantity (3), which measures the difference between the two transformed correlation coefficients, is approximately normally distributed with mean zero and variance one:

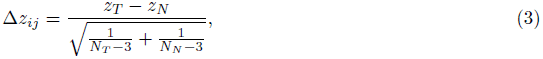

where *N*_*T*_ is the number of tumor samples, *N*_*N*_ is the number of normal samples, *z*_*T*_ is the Fisher-transformed tumor sample correlation coefficient, and *z*_*N*_ is the Fisher-transformed normal sample correlation coefficient. Thus, we can associate p-values 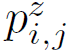 with the Z-test in (3) for each pair of genes *i, j*. After again correcting *p*^*z*^ for multiple hypothesis testing using the Bonferonni correction, we stored the results of our calculations in a matrix **D** with entries

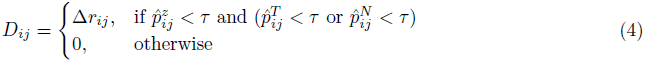

where 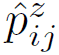 is the Bonferonni adjusted value of 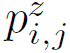. The entries of the matrix **D** correspond to the change in gene co-expression between tumor and normal samples, and will be our main object of study. We emphasize one final, but important, feature of Equation 4: an entry of **D** is nonzero if and only if that gene pair shows *both* (1) a significant change in co-expression between tumor and normal samples, *and* (2) the genes were co-expressed at a statistically significant level in tumor or normal samples (or both).This ensures that those gene pairs which we call differentially co-expressed are also co-expressed at a statistically significant level in at least one group of samples.

### Differential Co-Expression in Tumor vs. Normal Tissues is Not Random

With our analytical framework established, our first aim was to assess how pervasive differential co-expression was among metabolic genes in cancer samples. To do so, we applied the differential co-expression analysis described above to two TCGA studies (breast, BRCA; and clear cell renal cell carcinoma, KIRC) with large numbers of both tumor and normal RNA-Seq samples (106 and 71 normal samples, 914 and 480 primary tumor samples, respectively). We used a strict Bonferonni corrected p-value threshold of 1×10^*−*2^ to identify pairs of genes which we called differentially co-expressed. Across the total number of pairs of metabolic genes in our dataset (approximately (2×10^3^)^2^*/*2 = 2 × 10^6^ distinct pairs), we calculated (for each of the two studies) that approximately 2.5 percent of gene pairs were differentially co-expressed.

To independently test the extent of differential co-expression in our data, we followed the protocol presented in [2] and completed a permutation test to assess how frequently we would expect the observed changes in correlation coefficients by chance (Figure S1). In this analysis, we shuffled the labels (*e.g.* tumor or normal) of all samples, and calculated the difference in correlation coefficients and transformed correlation coefficients in the new, permuted data. This process was repeated 100 times, and the results aggregated to form a distribution. Visual inspection of the results confirmed that for a large number of gene pairs, the differences in correlation coefficients were larger in the real data than in the permuted data (Figure S1). Although it was computationally intractable to complete enough permutations of the data to generate robust p-values, we nevertheless found that 31% of gene pairs showed a higher difference in both (1) tumor and normal correlation coefficients and (2) transformed correlation coefficients than in any of the 100 permuted data sets. These findings supported our observation of extensive differential co-expression in metabolic genes.

Naturally, we were interested in identifying those genes which were enriched for membership in differentially co-expressed gene pairs. To find these genes, we calculated two “scores” for each gene:

1. 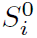, the number of differentially co-expressed gene pairs which gene *i* participates in
2. 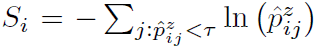, a weighted sum of the number of differentially co-expressed pairs gene *i* participates in

The score **S**, based on Fisher’s method for combining p-values from independent statistical tests [47], accounted for both the frequency of a gene’s membership in differentially co-expressed pairs, as well as the confidence with which we could claim the gene pair was differentially co-expressed (*i.e.* by the magnitude of 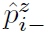). It is important to note that each test of differential co-expression in our dataset is not independent, so we cannot use **S** as a formal test statistic. However, its use as a measure of the recurrence and magnitude of a gene’s overall differential co-expression is nevertheless useful.

In addition to to the two metrics described above, we also wanted to make special note of those pairs of differentially co-expressed genes which took part in a known, previously reported biological interaction. To do so, we extracted from the Pathway Commons database [8] a list of pairs of genes known to interact in either of two ways: 1) through the formation of a complex with each other (“In-Complex-With” interactions), and 2) through the production of a metabolite by the enzyme encoded by one gene in the pair, and subsequent use of that metabolite as a substrate for the enzyme encoded by the other gene in the pair (“Catalysis-Precedes” interactions) [5]. We then identified which pairs of differentially co-expressed genes participated in either of these kinds of interactions. These results were summarized in two additional gene-level statistics, 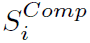 and 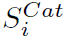, indicating the number of differentially co-expressed catalysis-precedes and in-complex-with interactions, respectively, a gene *i* participates in.

Our metrics highlighted a common feature of differential co-expression patterns in both BRCA and KIRC: differential co-expression was not randomly distributed throughout the metabolic network. Instead, most genes participated in relatively few differential co-expression interactions, while a small subset of “hub” genes participated in hundreds ( Figure 2C, histograms). In Tables 2 and 3, we report the top ten scoring genes (sorted by the metric 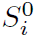) for the breast and kidney TCGA studies. The lists include a number of genes known to be associated with cancer, as well as a number of previously unreported genes. In the supplementary material, we provide a complete list of the scores for each metabolic gene in our study.

**Table 2.** Top differentially co-expressed genes in BRCA.

| Entrez | HUGO | $S_i$ | $S_i^0$ | $S_i^{\text{Cat}}$ | $S_i^{\text{Comp}}$ |
| --- | --- | --- | --- | --- | --- |
| 38 | ACAT1 | 11731.9 | 516 | 11 | 0 |
| 5264 | PHYH | 13080.2 | 512 | 3 | 0 |
| 10449 | ACAA2 | 10819.4 | 508 | 12 | 0 |
| 9588 | PRDX6 | 10983.9 | 506 | 2 | 0 |
| 4259 | MGST3 | 9211.37 | 494 | 1 | 0 |
| 7360 | UGP2 | 9882.05 | 492 | 0 | 0 |
| 847 | CAT | 9462.16 | 490 | 3 | 1 |
| 48 | ACO1 | 11005.9 | 484 | 3 | 0 |
| 7371 | UCK2 | 6024.4 | 483 | 0 | 0 |

**Table 3.** Top differentially co-expressed genes in KIRC.

| Entrez | HUGO | $S_i$ | $S_i^0$ | $S_i^{Cat}$ | $S_i^{Comp}$ |
| --- | --- | --- | --- | --- | --- |
| 29968 | PSAT1 | 13719.8 | 492 | 0 | 0 |
| 10846 | PDE10A | 11676.9 | 474 | 3 | 0 |
| 6482 | ST3GAL1 | 13591.9 | 466 | 0 | 0 |
| 1594 | CYP27B1 | 12064.4 | 465 | 0 | 0 |
| 65010 | SLC26A6 | 10902.9 | 451 | 0 | 0 |
| 64849 | SLC13A3 | 15016.4 | 445 | 0 | 0 |
| 64077 | LHPP | 7200.28 | 444 | 2 | 0 |
| 205 | AK4 | 9709.72 | 443 | 0 | 0 |
| 54682 | MANSC1 | 9081.81 | 440 | 0 | 0 |

In breast cancer, the top-ranked gene was ACAT1 (Acetyl-CoA acetyltransferase, not be confused with the enzyme acyl-Coenzyme A: cholesterol acyltransferase 1, which is encoded by the gene SOAT1). The enzyme translated from ACAT1 catalyzes the formation of acetoacetyl-COA, and along with acetyl-CoA is the precursor to 3-hydroxy-3-methylglutaryl-CoA. These two metabolites lie at the beginning of the mevalonate pathway, which generates precusors for cholesterol and steroid biosynthesis. Intriguingly, Freed-Pastor and colleagues [19] recently reported that upregulation of the mevalonate pathway is sufficient and necessary for mutant p53 to have phenotypic effects on cell architecture in mammary tissue. Overexpression of various genes in the mevalonate pathway has also been shown to associate with poor prognosis in breast cancer [13]. Interestingly, ACAT1 has a very high **S^Cat^**, indicating it is differentially coexpressed with 11 genes for which it is a catalytic partner: ACAA2, DLD, MLYCD, HADHB, HADH, OXCT1, PCCA, PDHA1, PDHB, and ACSS1. A plot of the differences in the correlation of these genes with ACAT1 is in Figure S4. In many cases, the co-expression patterns show remarkably tight correlations in normal tissue, and these correlations are partially or completely eroded in the tumor samples. Functionally, many of these genes are part of the terminal reactions in glycolysis, lipid biosynthesis and fatty acid oxidation. This loss of co-expression suggests that the flux generated by these pathways is no longer coupled to the flux through ACAT1 in tumor cells.

For KIRC, the highest-scoring differentially co-expressed gene was PSAT1 (phosphoserine aminotransferase 1), a key enzyme in the serine biosynthesis pathway which has already been associated with breast and colorectal cancers before [38, 45], but has not yet been associated with kidney cancer. PSAT1 was differentially co-expressed with 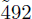 other metabolic genes in the dataset, with the strongest signals coming from genes like GATM (glycine aminotransferase), GBA3 (a beta-glucosidase), and SLC10A2 (a bile transporter) (Figure S2). Because nearly all of the strongest signals came from loss of positive correlation in normal samples, we further identified those genes with which PSAT1 was more strongly co-expressed in tumor samples qthan in normal samples (Figure S3). These genes included several galactosidases (GLA, GLB1), glycogen phosphorylase (PYGB), and SLC35A2, which transfers nucleotide sugars into the Golgi body for the purposes of glcosylation. Neither the substrates (3-phosphonoxypyruvate,2-oxoglutarate) nor the products (phosphoserine, glutamate) of PSAT1 participate in the glycogenolysis pathway, suggesting that the positive correlation between PSAT1 and glycogen breakdown in tumors may be the result of indirect couplings. In particular, it is possible that the overexpression of glycogen phosphorylase may liberate carbon units to be shunted from glycolysis into the serine biosynthesis pathway through PSAT1, as well as into the Golgi body for glycosylation in tumor cells.

Following our analysis of PSAT1, we reasoned that a particularly interesting set of genes were those showing a higher degree of co-expression (as quantified by the magnitude of the Spearman correlation coefficient) in tumor samples relative to normal samples. For both BRCA and KIRC, we isolated pairs of genes exhibiting this property, and scored each metabolic gene based on how many such interactions it participated in. Interestingly, in both studies the highest-scoring gene was associated with the metabolism of lipids. In KIRC, the highest scoring gene was mevalonate kinase, MVK, a key gene in the cholesterol pathway described above for BRCA. In breast tissue, the highest scoring gene was LIPG, an endothelial lipase which catalyzes the hydrolysis of lipids. The products of this hydrolysis can then be used for the production of signaling lipids as well as cell membrane components.

### Breast and Kidney Cancers Show Distinct Co-Expression Patterns

Surprisingly, breast and kidney cancers shared no common genes among the ten highest-scoring genes in the differential co-expression analysis above. Prompted by this observation, we decided to investigate more explicitly whether differential co-expression patterns were similar between the two cancer types.

We reasoned that while individual genes may not show similar patterns of differential co-expression, larger groups (*e.g.* pathways) of genes might. To probe whether common, “global” patterns of differential co-expression existed between the two studies, we completed a principal components analysis (PCA, Figure 2A). We assembled a concatenated differential co-expression matrix:

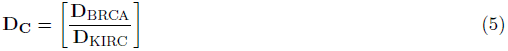

with dimension 2*m* × *m*, where *m* is the number of genes under study. For a given index *i < m*, row *i* corresponded to the differential co-expression pattern of that gene in BRCA, while row *m*+*i* corresponded to the differential co-expression pattern in KIRC. Thus, each column of **D_C_** corresponded to a metabolic gene, and stored the differential co-expression of that gene with all other metabolic genes in both breast and kidney studies.

Our expectation was that PCA would identify patterns of differential co-expression which breast and kidney cancers might share in common. Instead, we found that genes in the two studies displayed completely distinct patterns of differential co-expression ( Figure 2A). While a large portion of the variance in the data was captured by the first two principal components (33 and 19 percent of the total variance in the data, respectively), most genes from breast cancer had nearly no loading on component 2, while most genes from kidney cancer had nearly no loading on component 1. The result was the cross pattern evident in Figure 2A.

These discrepenacies between tumor types led us to directly compare the frequency of differential co-expression for each gene in the two studies ( Figure 2C). We observed a remarkably high density of points which fell on either of the two axes, corresponding to genes which showed no differential co-expression in one of the tumor types. Furthermore, of the 1789 genes we examined, we identified 224 genes (13% of all genes) which participated in at least 10 differential co-expression pairs in breast cancer, but less than 2 in kidney cancer, and 164 genes with at least 10 differential co-expression pairs in kidney cancer, but less than 2 in breast cancer.

Of particular interest to us were genes which showed extreme cases of the pattern described above: very high differential co-expression in one tumor type, but none in the other. To identify these genes, we calculated the mean and standard deviation of **S^0^** (the number of differentially co-expressed gene pairs a gene participates in) for each study. We then searched for genes with **S^0^** greater than two standard deviations above the mean **S^0^** in one study, but with **S^0^** = 0 in the other study ( Figure 2C, blue and green points). Doing so, we identified dozens of genes which seemed to be specifically de-coupled in only one of the two studies. Among these, the kidney specific genes seemed to be highly enriched for SLC and ABC transporters. A particularly interesting kidney-specific gene was DPEP1 (a dipeptidase) in light of the recently observation of elevated dipeptide levels in a subset of clear cell renal carcinoma tumors (manuscript in preparation). In contrast, breast-specific genes included CDO1 (cysteine dioxygenase Type 1, whose inactivation was recently reported to contribute to survival and drug resistance in breast cancer [27]) and a number of genes involved in glycerolipid/lipid biosynthesis and associated with malignancy in breast cancer (GPAM [15] and MGLL [37]).

We were similarly interested in finding genes enriched for differential co-expression in both BRCA and KIRC samples. Among them was ASS1, an enzyme which catalyzes the rate-limiting step in arginine synthesis, and is further invovled with the synthesis of nitric oxide and polyamines. Perhaps the most intriguing feature of arginine metabolism in cancer is that several tumor types exhibit an arginine auxotrophy phenotype, and are unable to proliferate in the absence of arginine [14]. Intriguingly, Qiu and colleagues recently reported the killing of triple-negative breast cancer cell lines under arginine deprivation, identifying it as a lucrative therapeutic target [39]. It is not clear from our analysis whether differential co-expression of ASS1 is associated with such a vulnerability, but its recurrent differential co-expression in both studies suggests that its activity may play an important role in malignancy.

Finally, we analyzed the pattern of differential co-expression across metabolic pathways, as annotated in the Recon2 metabolic network [43] (see Methods). The results of our analysis are highlighted in Figure 3B, where we compared the score of each pathway in BRCA and KIRC, respectively. In breast cancer, among the most enriched pathways is peroxisomal transport genes, including the peroxisomal transporters ABCD1, ABCD2, and ABCD3, which transport fatty acids and acyl-CoAs and have been shown to be markers of tumor progression and response to therapy [22]. Notably, genes in the vitamin C pathway were enriched for differential co-expression in both cancers, possibly as an indirect consequence of high oxidative stress within the tumors.

**Figure 3.**
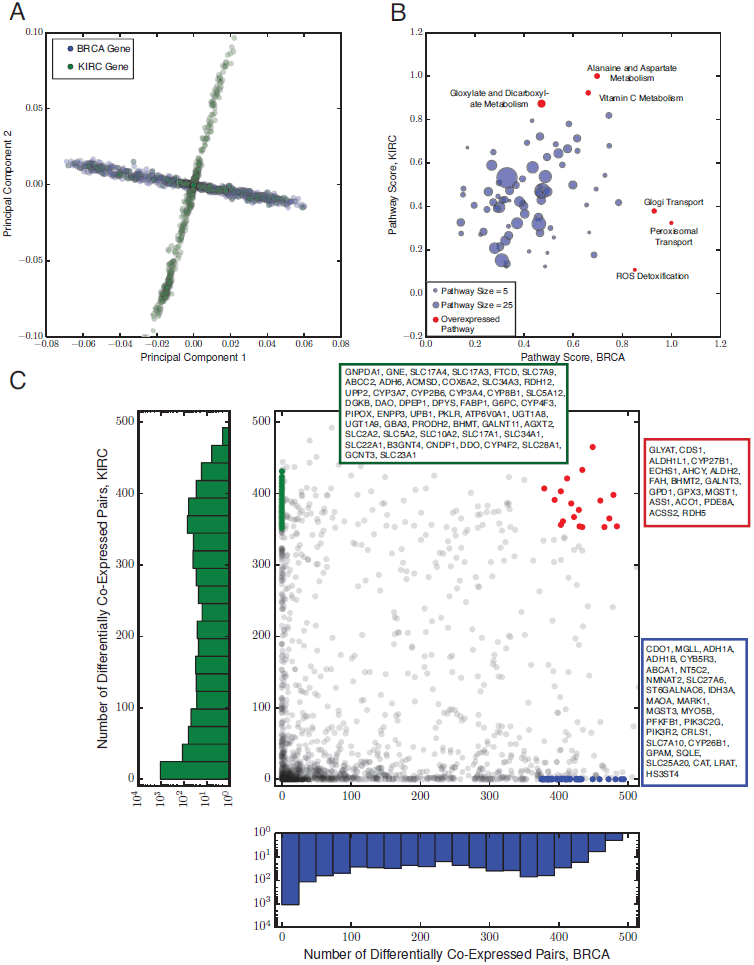
(**A**) Principal components analysis (PCA) for the breast (green) and kidney (blue) differential co-expression data. Each dot represents one gene. Data from kidney tumors exhibits variation mostly along the first principal component, while data from breast tumors varies mostly along the second, suggesting that the dominant modes of variation in the two tumor types are distinct from each other. (**B**) Differential co-expression pathway analysis. Each axis denotes the enrichment score for a pathway in breast or kidney tumors, respectively. Red dots indicate significantly over-or under-enriched pathways. (**C**) A comparison of the score 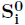 in breast and kidney tumors. Each dot is a single gene. A number of genes (blue and green dots and inset boxes) show extensive differential co-expression in one tumor type, but none in the other. Other genes (red dots and inset box) are highly differentially co-expressed in both.

### Differential Expression Sheds Little Light On Differential Co-Expression

So far, we have spent a great deal of effort identifying individual genes which seem to be co-expressed in different ways in normal and tumor tissues. A more common first step in the analysis of gene expression data across samples is the identification of differentially expressed transcripts. The underlying rationale behind differential expression analysis of metabolic genes is intuitive: that higher expression of genes in one condition over another suggests a difference in the metabolic flux through those sets of genes. In this study, we are more concerned with the coupling of genes together: since metabolic genes are components of a network, differential regulatory patterns (which may not necessarily substantially affect absolute expression levels) may lead to differences in metabolic flux. Naturally, one may ask whether the two sets of genes overlap; in other words, do genes which are up- or down-regulated in tumor (compared to normal) also exhibit large differences in co-expression patterns in tumor (compared to normal) samples? At first glance, one might expect the answer to be yes. The coordinated over-/under-expression of genes may seem advantageous to the cell: if the cell’s objective is to increase the flux through a metabolic pathway, it makes sense to synchronously overexpress each constituent enzyme of that pathway. However, it is well-known that metabolic pathways route flux in non-canonical ways, especially in cancer (*e.g.* aerobic glycolysis, overflow metabolism of 3-phosphogylcerate into one-carbon metabolism [48]).

To explicitly test the connection between differential co-expression and differential expression, we compared the two measures for metabolic genes in BRCA and KIRC ( Figure 3). We assessed differential expression using the limma voom package [30]. We found that the magnitude of differential expression (as quantified by the log_2_ ratio of tumor to normal expression) was weakly associated with the frequency of differential co-expression of a gene (BRCA, Spearman *ρ* 0.21; KIRC, Spearman *ρ* 0.11). In spite of this weak association, many of the most differentially expressed genes were members of very few dysregulated gene pairs, and conversely many genes which exhibited no substantial change in expression levels nevertheless were found to be frequent members of dysregulated gene pairs.

The most intriguing observation we made was that a number of genes showed no measurable change in absolute expression levels, but nevertheless were among the most differentially co-expressed genes in the entire dataset (green dots, Figure 3). To find exceptional cases like these, we identified genes with **S^0^**greater than 2 standard deviations above the mean **S^0^** for the study, but with an absolute log_2_ ratio of less than 0.2. For breast cancer, these genes included PLOD2 (procollagen lysyl hydroxylase 2 [20], recently reported to be essential for hypoxia-induced breast cancer metastasis), and LDHA, a key enzyme in the terminal end of glycolysis. In KIRC, several of the genes we identified (RENBP, GNE, and CTSA) were members of the glycoprotein sialyation pathway, which has also been associated with metastasis [1].

The presence of genes with exceptionally high differential co-expression and eseentially no differential expression (and the converse) deserves further discussion. It is possible that, depending on how the activity of a metabolic pathway is modulated, either differential expression or differential co-expression may be a more suitable technique for identifying such modulation. In one case, a gene may change in synchrony with its regulatory partners; that is, regardless of whether the gene is over- or under-expressed relative to normal tissue, it exhibits precisely the same co-expression patterns. Such an effect may be observed, for example, following the over-expression of a transcription factor common to all the genes in a co-expressed cluster. As we suggested earlier, synchronous regulation of a metabolic pathway may serve as a mechanism for increasing flux through the pathway, and would be detected through standard differential expression analysis. In contrast, a gene’s expression may correlate with different sets of genes in different conditions. In our case, the control over expression wielded by one transcription factor in normal tissue *TF*_*N*_ would be ceded to a different transcription factor in tumor tissue *TF*_*T*_. The consequence is that the gene of interest is co-expressed with a completely distinct set of genes under the control of *TF*_*T*_. The differential co-expression of such a gene provides indirect evidence that the source or destination of metabolic flux through the enzyme encoded by this gene may be changing from normal to tumor tisues.

### Signatures of Regulation in Differential Co-Expression Patterns

As alluded to above, the expression of genes is fundamentally orchestrated by regulatory factors such as transcription factors and microRNAs. Thus, the differential co-expression patterns we observe are likely due, at least in part, to differential regulatory activity by these molecules. Inspired by prior work linking transcription factors with observations of differential co-expression [24, 40], we examined our differential co-expression networks for an enrichment of targets associated with particular transcription factors annotated in MSigDB [32]. To detect such enrichment, we isolated metabolic genes which were reported targets of a particular transcription factor. Then, we applied a binomial test (see Methods) to quantitatively assess whether the number of differential co-expression edges existing between only these target genes was higher than would be expected by chance. We used only highly significant differentially co-expressed edges, with a p-value threshold of 1×10^*−*4^.

Among the 200 or so transcription factors we examined, only a few dozen were enriched in either kidney or breast cancer. In breast cancer, the most enriched transcription factors (reported in Table 4) included SP1, NFAT, and ERR1. Several of these transcription factors have already been reported to play important roles in breast cancer throughout the literature. SP1 is known to be involved in cell proliferation, apoptosis, and cell differentiation and transformation, and has been reported as a prognostic marker for breast cancer [11, 31]. Both NFAT and SP1 have been shown to induce invasion of breast tissue via the transcriptional modulation of downstream genes [11, 49]. Perhaps most interesting is the identification of ERR1 (estrogen-related-receptor 1, also known as as ERR-*α*), an orphan receptor known to interact with PGC1-*α* to regulate a number of metabolism-related genes. ERR-*α* is regulated by ErbB2/Her2 signaling [4], and is associated with poor outcomes in breast cancer patients [3].

**Table 4.** Transcription factors most enriched for differential co-expression targets in BRCA.

| MSigDB Transcription Factor Motif | P-Value |
| --- | --- |
| SP1-Q6 | $2 \times 10^{-22}$ |
| HNF4-01 | $9 \times 10^{-16}$ |
| NFAT-Q4-01 | $3 \times 10^{-13}$ |
| FREAC2-01 | $1 \times 10^{-11}$ |
| ETS2-B | $2 \times 10^{-8}$ |
| AACTTT-UNKNOWN | $6 \times 10^{-8}$ |
| ERR1-Q2 | $8 \times 10^{-8}$ |
| YNGTTNNNATT-UNKNOWN | $5 \times 10^{-7}$ |
| HFH8-01 | $8 \times 10^{-7}$ |
| CTTTAAR-UNKNOWN | $4 \times 10^{-6}$ |

For kidney cancer, the pattern was far more unanimous: several of the most enriched transcription factor target sites were targets of HNF4 (Table 5). HNF4 is known to control cell proliferation in kidney cancer cell lines, and regulates a number of well-known cancer-associated genes to do so (*e.g.* CDKN1A and TGFA) [21, 35, 42]. Interestingly, HNF4A (one of the two isomers of HNF4, which was most enriched for differential co-expression targets) shows no clear differential expression pattern between KIRC tumor samples and adjacent normal tissue samples ( Figure 5A), but does seem to exhibit more variation in tumor samples than in normal samples. The co-expression of HNF4 and its metabolic gene targets is markedly different in normal and tumor samples ( Figure 5B). A number of these genes (including PIK3R3, a member of the PI3K pathway, and PKLR, an isoform of pyruvate kinase) showed exceptionally strong co-expression with HNF4A in normal samples, only to have this co-expression abrogated in tumor samples (Figure S5C). Similarly, many of the strong co-expression patterns existent *between* the targets of HNF4A and HNF4A itself in normal samples wre also abrogated in tumor samples ( Figure 5B). Together, these findings suggest that the regulatory program associated with HNF4A in normal tissue is disrupted in tumor tissue, a hypothesis in line with previous findings implicating its dysregulation with increased cell proliferation [21]. Given its high score in our enrichment analysis, we tested whether the expression of HNF4A was associated with patient survival in the TCGA data. After stratifying patients into groups with high and low expression (relative to the mean expression of HNF4A in the tumor samples), we found that low HNF4A expression is associated with shorter survival in KIRC patients ( Figure 5D, log-rank p-value 0.007).

**Figure 4.**
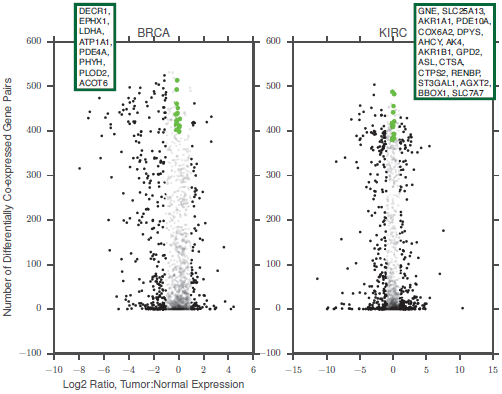
Differential expression and differential co-expression are only weakly correlated. Green dots, detailed in insets, indicate genes with high differential co-expression score 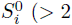 standard deviations above the mean 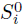) but very small absolute fold ratio (*<* 0.1). Black dots indicate a gene is differentially expressed with corrected p-value less than 0.01 and absolute fold log2 fold ratio greater than 1. Transparent dots correspond to genes which are not differentially expressed.

**Figure 5.**
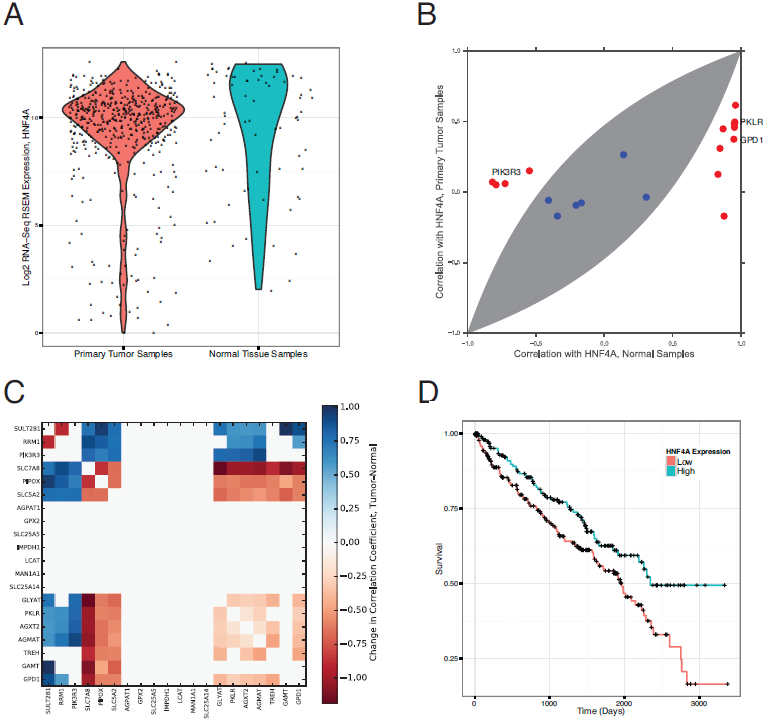
The targets of HNF4 are enriched for differential co-expression in KIRC. **(A)** HNF4A is not differentially expressed between tumor and adjacent normal tissue samples in KIRC. Each dot corresponds to the expression of HNF4A in one sample of either primary KIRC tumor or normal kidney tissue. (**B**) Nevertheless, the metabolic gene targets of HNF4A show a distinct loss of co-expression with HNF4A in tumor samples. Several of these genes reside in central carbon metabolism. Genes outside the shaded area correspond to statistically significant instances of differential co-expression. (**C**) Heatmap of differential co-expression for the 20 metabolic gene targets of HNF4A containing the motif AARGTCCAN around the transcription start site. Value of each square indicates the difference in correlation coefficients between tumor and normal samples, with statistically insignificant differences set to zero. A strict p-value threshold of 1 × 10*−*4 was used to assign statistical significance. **(D)** Survival curves for patients showing low or high expression of HNF4A. Patients with low expression of HNF4A exhibited worse outcomes.

**Table 5.** Transcription factors most enriched for differential co-expression targets in KIRC.

| MSigDB Transcription Factor Motif | P-Value |
| --- | --- |
| LEF1-Q2 | $4 \times 10^{-45}$ |
| HNF4-Q6 | $2 \times 10^{-31}$ |
| HNF4-01 | $7 \times 10^{-30}$ |
| HNF4ALPHA-Q6 | $7 \times 10^{-24}$ |
| HNF4-DR1-Q3 | $1 \times 10^{-13}$ |
| HNF4-01-B | $4 \times 10^{-10}$ |
| COUP-01 | $4 \times 10^{-8}$ |
| PPAR-DR1-Q2 | $6 \times 10^{-7}$ |
| PAX4-02 | $6 \times 10^{-5}$ |
| IK3-01 | 0.0002 |

Taken together, our observations above suggest that HNF4A’s control over the expression of its targets changes in at least a subset of clear cell kidney tumors when compared to normal kidney tissue. It is possible that this loss of control occurs via under-expression of HNF4A itself. It is also possible that (as we proposed in the prior section) other transcription factors exert a more dominant control over HNF4A’s targets. In either case, this leads to the loss of co-expression among HNF4A’s targets, and between HNF4A itself and its targets.

### PanCan Patterns of Differential Co-Expression

This final section of our work strikes out into more difficult territory: we ask whether some patterns of differential co-expression may exist throughout different cancer types, regardless of their tissue of origin. While we have found a number of apparently dysregulated metabolic genes specific (and in some cases, common) to breast and clear cell renal cell carcinoma tumors, we have made little effort to search for common patterns across many different types of tumors. Such a search is necessarily complicated by the fact that our analytical method requires large numbers of normal and tumor samples for sufficient statistical power. The TCGA features few studies with large numbers of normal RNA-Seq samples. In order to balance the need for statistical power with our desire to detect so-called “PanCan” patterns of differential co-expression, we included five more studies (lung adenocarcinoma, LUAD; hepatocellular carcinoma, LIHC; prostate adenocarcinoma, PRAD; head and neck squamous cell cancer, HNSC; and thyroid cancer, THCA) with at least 30 normal RNA-Seq samples, in our analysis. To increase the confidence of our predictions, we used a stricter p-value threshold of *τ* = 1 × 10^*−*4^ to call statistically significant differential co-expression. The results of the PanCan analysis are shown in Figure 6. We retained only those genes which were members of a gene pair which was differentially co-expressed in at least three of the seven studies. Out of the 1789 metabolic genes under study, only 50 genes satisfied this criteria. Interestingly, many of these genes encode key enzymes in central metabolism (for example, PC, pyruvate carboxylase; LDHD, D-lactate dehydrogenase; IDH1, isocitrate dehydrogenase 1; ALDOA, aldolase A), pointing to apparently recurrent dysregulations of core pathways.

**Figure 6.**
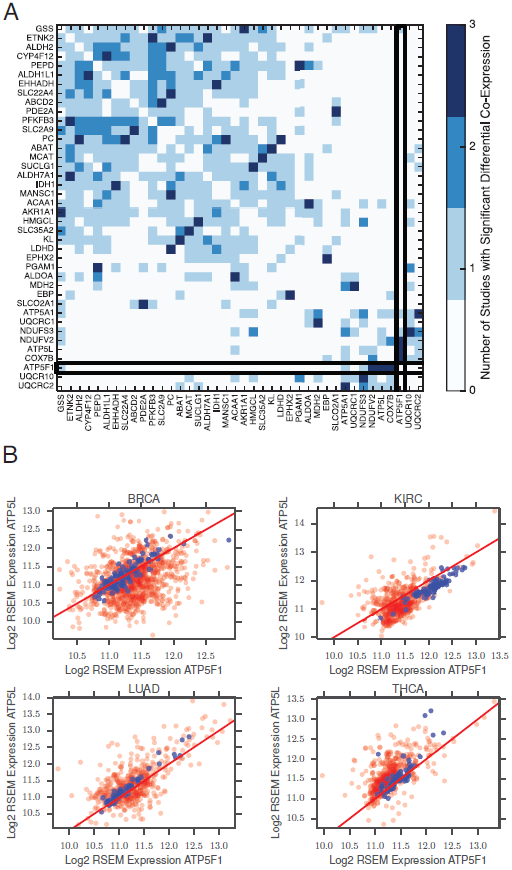
PanCan analysis of differential co-expression. **(A)** All gene pairs which showed differential co-expression in at least 3 out of 4 different TCGA studies were identified. Approximately 50 unique metabolic genes participated in these recurrently differentially co-expressed pairs. The differential co-expression across all possible pairs of thes genes is depicted in the heatmap. A p-value threshold of 1 × 10^*−*4^ was used to assign statistical significance for differential co-expression. Special emphasis is placed on ATP5F1. **(B)** Co-expression of ATP5F1 and ATP5L, both members of mitochondrial Complex V, in four different TCGA studies (blue dots: normal tissue samples; red dots: tumor samples). Red line corresponds to perfect 1:1 correlation. Tumor samples exhibit substantially noisier co-expression of these two genes.

Among the many individual results of our PanCan analysis, perhaps the most interesting was the recurrent dysregulation of four genes in the mitochondrial electron transport chain (ETC): two genes associated with mitochondrial ATP synthase complex V (ATP5F1 and ATP5L), COX7B (part of the complex IV cytochrome c oxidase), and NDUFV2 (complex I). A number of other mitochondrial ETC genes are also differentially co-expressed (but to a lesser extent), including UQCR10, UQCRC2, UQCRC1, ATP5A1, and NDUFS3. Given how critical these protein complexes are to energy production and proliferation, we examined in detail the co-expression patterns of ATP5F1 and ATP5L. We found an exceptionally strong correlation in the expression of both genes in normal tissue. Across all seven studies, the expression of both genes was almost precisely equal ( Figure 6B, blue dots). However, in tumor samples, the strength of the co-expression (as measured by the correlation coefficient) was substantially weaker. Notably, ATP5F1 and ATP5L were not differentially expressed; instead, their co-expression simply appeared “noisier” in tumor samples. To quantify whether this “noisier” co-expression may be occuring by chance, we fit each co-expression pattern in Figure 6B to a line, and then calculated the variance of the residuals of the fit. We used Levene’s test to test whether the variance of the residuals associated with tumor samples was larger than the variance of the residuals associated with normal samples. In all four tumor types, we confirmed that the tumor samples showed higher variance (p-value 7 × 10^*−*17^, 3 × 10^*−*3^, 6 × 10^*−*9^, 6 × 10^*−*6^, 3 × 10^*−*11^, 2 × 10^*−*3^, 5 × 10^*−*3^ for BRCA, HNSC, KIRC, LIHC, LUAD, PRAD, and THAC samples, respectively).

The functional consequences of these increasingly “noisy” co-expression patterns in ATP5F1 and ATP5L are unclear. It is known that stoichiometric imbalances of proteins (for example as a result of changes in gene dosage) in complex with each other can manifest phenotypically [44]. Given the recurrence of differential co-expression of three different gene pairs containing ATP5F1 and a second mitochondrial matrix member (ATP5L, COX7B, and NDUFV2), it is tempting to speculate that differential co-expression of ATP5F1 may lead to an altered mitochondrial phenotype. In particular, an imbalance in the levels of ATP5F1 and ATP5L may cause defects in the ability of mitochondria to efficiently conduct oxidative phosphorylation via the electron transport chain. Further experiments are required to evaluate this hypothesis.

## Discussion

In this work, we have searched for signals of differential co-expression in tumors. Among our findings, the most relevant is simply the prevalence of differential co-expression throughout metabolism. Gene expression studies are frequently the “first-step” analytical method of choice for understanding the consequences of a perturbation on an organism, or for the comparison of two distinct subsets of samples. While standard methods for differential expression analysis offer useful insights into the differential regulation of genes, our findings here (and the prior findings of others studying differential co-expression) suggest that a great deal of information remains to be culled from the study of “second-order” co-expression patterns between pairs of genes. We have shown that these two measures (differential expression and differential co-expression) are not interchangeable, and in many cases point to distinct regions of the metabolic network that may be dysregulated. Of course, it is important to remember that while the statistical power of both approaches relies on large sample sizes, differential co-expression is significantly more sensitive to sample size upon multiple hypothesis correction because of the large number of independent statistical tests (equal to the square of the number of genes) under evaluation.

Our findings here are a small, first step in applying such a second-order analysis to cancer data, and in particular to the study of cancer metabolism. We have made a number of assumptions in order to make progress in the analysis, and these assumptions should be re-visited in future work. In particular, we have repeatedly assumed that the expression of a gene roughly correlates with the abundance of its translated protein product, and that this abundance correlates with enzyme activity. An entire field of theoretical study (metabolic control analysis, [18]) and a number of experimental studies (*e.g.* [12]) have shown that metabolite abundances are equally, if not more, important for the control of fluxes. We note, given an adequately large number of samples, an analogous “differential correlation analysis” is possible for metabolomics data. It would be especially interesting to compare the results from such an analysis with the analogous results using expression data.

One major concern with our results are the confounding effects of (1) contamination by stromal and immune cells, and (2) existence of heterogeneous tumor subtypes in the data. Tumor samples are often contaminated with mixtures of normal adjacent tissue and immune cells. Deconvolving the contribution of non-cancerous cells from the total signal obtained from a tumor sample remains a major computational challenge, and it is unclear how the contribution of this non-cancerous signal affects our differential co-expression results. A separate but related concern is the existence of distinct molecular subtypes in a set of samples (*e.g. ER*^+^, *ER*^*−*^ breast cancer samples). We have not made any efforts to tease apart the confounding effects of these distinct subtypes in our work. Interestingly, it possible that a significant portion of the differential co-expression signal we identify derives directly form these subtypes; in other words, the primary differences between subtypes may lie among the differentially co-expressed genes. Evaluating such a hypothesis will require substantially larger sample sizes. Nevertheless, we feel that a more careful analysis of such patterns after subtype separation and stromal deconvolution is a lucrative route for future studies.

Finally, we would like to comment on the complementarity of differential expression and co-expression which we have proposed. In the course of responding to environmental stresses and stresses, it is inevitable that some genes will be both differentially expressed as well as differentially co-expressed. We are not arguing that one measure is superior to the other; rather, each offers a different glimpse onto the response of a highly-connected network to a perturbation. Neither the over-expression of a single gene, nor an increase in the co-expression of a pair of genes, signals a change in a pathway’s activity. However, by monitoring both measures, one univariate and the other multivariate, one may obtain a more complete picture of the complex system under examination.

## Methods

### Data

All TCGA expression data were accessed using the Broad Institute Firehose, using the latest version as of Sep 21, 2013.

### Pathway Scores for Differential Co-Expression Breast and Kidney Cancers

We assigned each gene in our study to one or more pathways using the subsystem assignments in the Recon2 human metabolic reconstruction [43] (see Methods). Then, for each TCGA study, we calculated a score for each pathway *i*, *E*_*i*_, using:

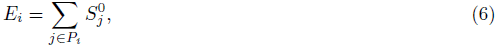

where **P_i_** is the set of all genes in pathway *i* and *m* is the total number of genes. Thus, *E_i_* counts the total number of dysregulations for all genes in pathway *i*. We then divided each *E*_*i*_ by the number of genes in pathway *i* to obtain a normalized pathway score 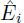. Thus, 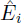 quantifies the differential co-expression of all genes in a pathway, averaged over the number of genes in that pathway. We excluded from our analysis pathways composed of fewer than five genes.

### Tests for enrichment of transcription factor targets

We obtained data on transcription factor targets from the Broad Institute’s MSigDB website [32]. TAssuming that a particular regulatory factor has *m* targets, we calculate the total number of differential co-expression edges in the sub-network composed of only these *m* gene targets. In this subnetwork, there are 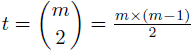 total possible edges. If we see *e* edges in the true subnetwork, we can calculate the probability that these edges would appear by chance. Given that the probability of a random edge in the network is *p* (for a Bonferonni-corrected p-value threshold of 1×10^*−*2^, *p ≈* 8×10^*−*3^), the probability of seeing at least *e* edges is

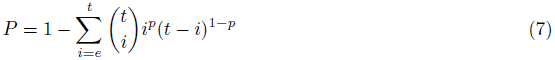

We then Bonferonni-corrected the p-value obtained from the calculation above.

## Supplemental Material

**Figure S1.**
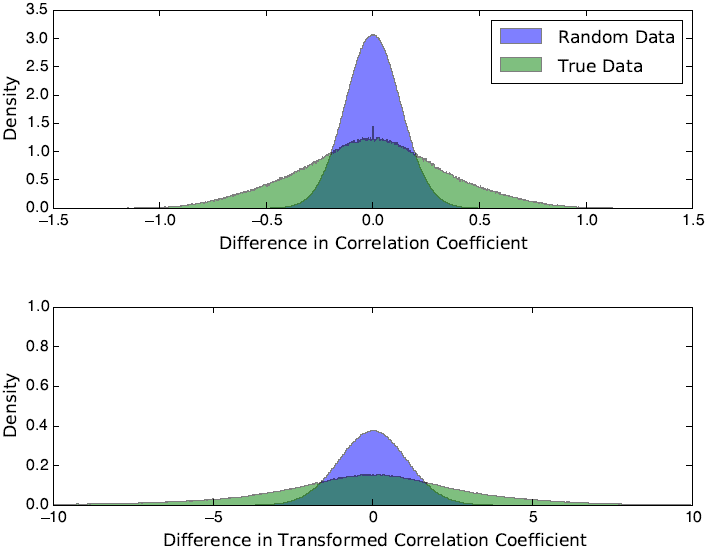
A comparison of the difference in correlation coefficient in clear cell kidney cancer for the true (green) and permuted random (purple) data sets. The labels (i.e. tumor or normal) of all RNA-Seq samples were permuted, and the difference in correlation coefficient calculated. This process was repeated 100 times to generate a distribution. Differences in correlation coefficients tend to be larger in the true data, suggesting that differential co-expression is being observed.

**Figure S2.**
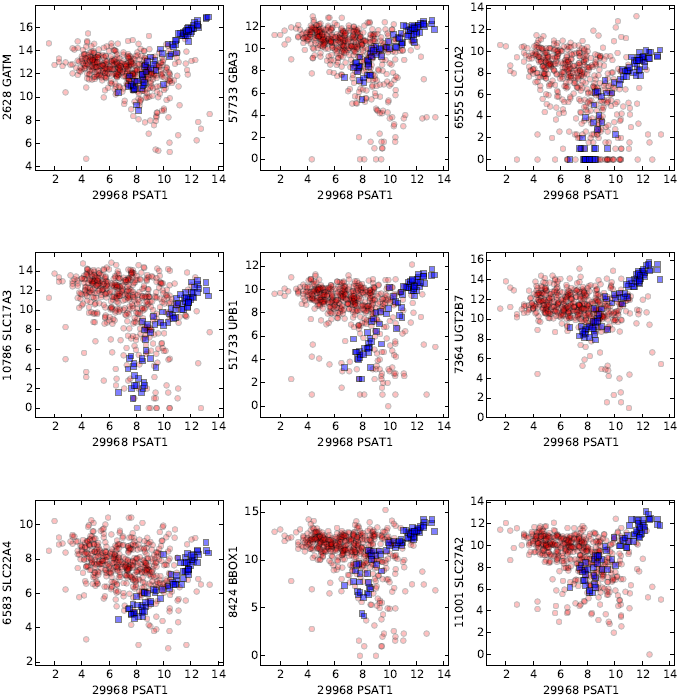
Differential co-expression in PSAT1 in KIRC.

**Figure S3.**
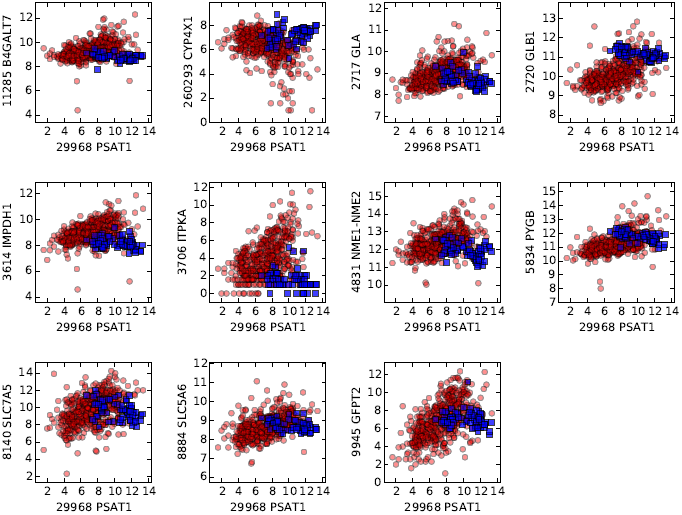
Novel co-expression in KIRC tumors with PSAT1.

**Figure S4.**
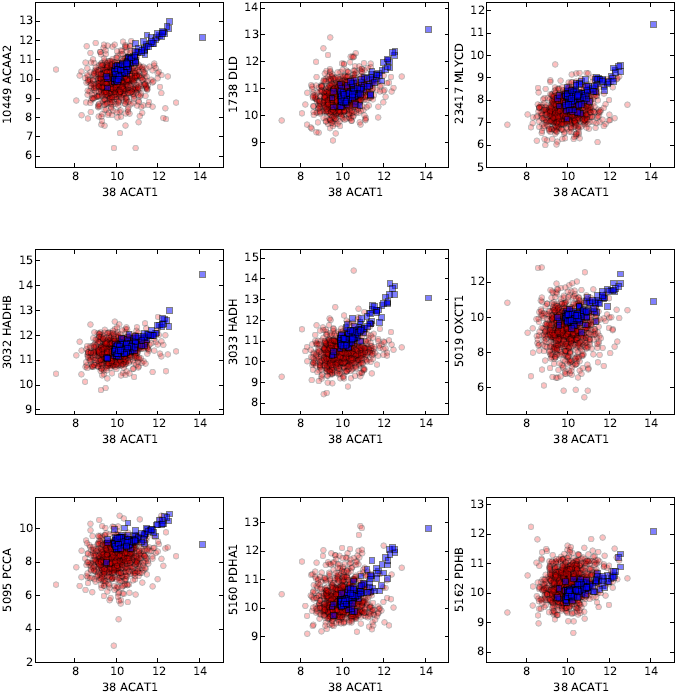
Differential co-expression in ACAT1 in BRCA.

**Figure S5.**
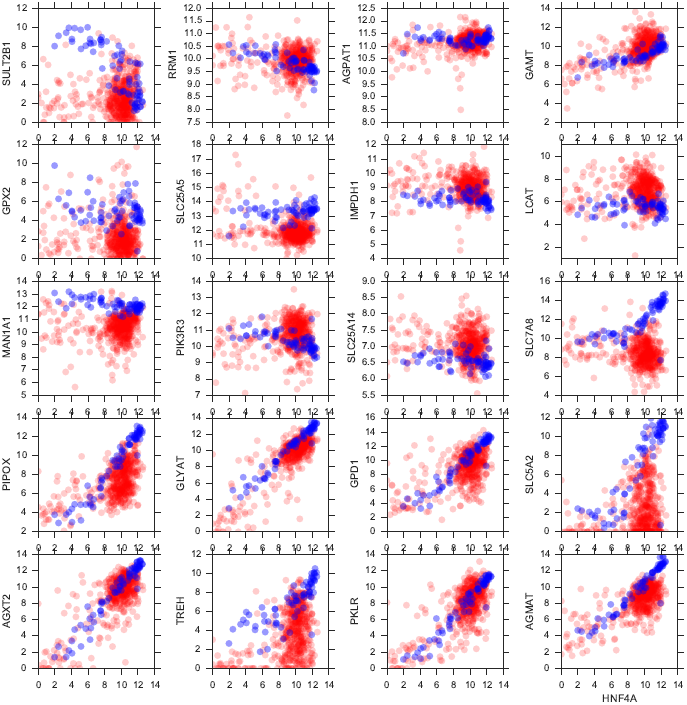
Differential co-expression of HNF4 with its metabolic gene targets in KIRC.

